# *Sclerotinia sclerotiorum* hijacks copper from its host for infection

**DOI:** 10.1101/2020.01.09.900217

**Authors:** Yijuan Ding, Jiaqin Mei, Yaru Chai, Wenjing Yang, Yi Mao, Baoqin Yan, Yang Yu, Joseph Onwusemu Disi, Kusum Rana, Jiana Li, Wei Qian

## Abstract

*Sclerotinia sclerotiorum* induces host reactive oxygen species (ROS) production, which leads to necrosis in the host, allowing the pathogen to absorb nutrients from the dead tissues. Here, we found that three *S. sclerotiorum* genes involved in copper ion import/transport, *SsCTR1*, *SsCCS* and *SsATX1*, were significantly up-regulated during infection of *Brassica oleracea*. Function analysis revealed that these genes involved in fungal ROS detoxification, oxalic acid production, pathogen establishment and virulence. On the host side, four genes putatively involved in copper ion homeostasis, *BolCCS*, *BolCCH*, *BolMT2A* and *BolDRT112*, were significantly down-regulated in susceptible *B. oleracea*, but stably expressed in resistant *B. oleracea* during infection. Their homologs were found to promote resistance to necrotrophic pathogens and increase antioxidant activity in *Arabidopsis thaliana*. Furthermore, copper concentration analysis indicated that copper is transported into the necrotic area from healthy area during infection. Collectively, our data suggest that *S. sclerotiorum* hijacks host copper to detoxify ROS, whereas the resistant hosts restrict the supply of essential copper nutrients to *S. sclerotiorum* by maintaining copper ion homeostasis during infection.

## Introduction

Copper serves as a cofactor in many enzymes and is an essential micronutrient for growth and development of organisms. It is involved in a range of biological processes, including photosynthetic and respiratory electron transport, cell wall remodeling, oxidative stress responses, and ethylene perception (Pilon *et al*, 2006; Yruela, 2009). Given its importance, copper metabolism has been well-studied in model organisms. In yeast, Cu^2+^ is reduced to Cu^+^ by cell membrane metalloreductases (Fre1 and Fre2), and Cu^+^ is then transported into cells by the high-affinity Cu^+^ transporters Ctr1 and Ctr3 (Pena *et al*, 2000). The Ctr2 transporter mobilizes the stored copper from the vacuole into the cytosol under low-copper conditions (Rees *et al*, 2004). The cytosolic copper is delivered to the cuproenzymes in diverse ways. For example, the copper homeostasis factor Atx1 binds and delivers copper into Fet3 via the Ccc2 pump in yeast (Lin *et al*, 1997; Cankorur-Cetinkaya *et al*, 2016). Copper chaperone CCS delivers copper into Cu/Zn SOD in human and yeast (Banci *et al*, 2012; Gleason *et al*, 2014), and copper is transferred into cytochrome *c* oxidase in the mitochondria via copper chaperones such as COX17 and COX11 in eukaryotes (Carr and Winge, 2003; Arnesano *et al*, 2005). A few genes related to the absorption and distribution of copper have been discovered in *Arabidopsis thaliana*, such as genes encoding copper transporters (COPTs), chaperone components (CCH, CCS and COX), metallothioneins (MTs), P-type ATPases (HMA, PAA and RAN) and plastocyanin (PETE) (del Pozo *et al*, 2010; Gu *et al*, 2015; Abdel-Ghany, 2009).

Reactive oxygen species (ROS) including hydrogen peroxide (H_2_O_2_), hydroxyl radical (HO-), singlet oxygen (^1^O_2_) and superoxide anion (·O_2_^-^), are derived from partial reduction of oxygen (O_2_) (Liu and He, 2017). ROS have been called ‘double-edged swords of life’ (Mittler, 2017). On the one hand, ROS act as signaling molecules that regulate development, differentiation, redox levels, stress signaling, interactions with other organisms and systemic responses (Mittler *et al*, 2011). On the other hand, excess ROS cause oxidative cellular injury to DNA, RNA, proteins and lipids, and also trigger programmed cell death (Mittler, 2017; Foyer and Noctor, 2013; Mignolet-Spruyt *et al*, 2016). To avoid or overcome the damage caused by excess of ROS, organisms have developed a complex ROS scavenging system that delicately regulates the balance between production and elimination of ROS. A few cuproenzymes are involved in ROS scavenging and antioxidant activity. For example, the cytosolic Cu/Zn superoxide dismutase (Cu/Zn SOD) constitutes the front-line defense against intra- and extracellular ROS (Culotta *et al*, 2006). Copper homeostasis factor ATX1 is involved in defense against oxidative stress (Himelblau *et al*, 1998), while cytochrome *c* oxidase catalyzes the reduction of oxygen to water in mitochondria (Poyton *et al*, 1995).

Necrotrophic plant pathogens promote ROS production in the plant host and induce necrosis during host colonization (Heller and Tudzynski, 2011). This raises an interesting question of how necrotrophic plant pathogens survive in such high levels of host-derived ROS. *Sclerotinia sclerotiorum* is a typical necrotrophic pathogen that causes *Sclerotinia* stem rot in more than 400 species (Garg *et al*, 2010). In this study, our data showed that *S. sclerotiorum* hijacks copper from the host and activates ROS detoxification enzymes during infection by enhancing the expression of genes involved in copper ion import and transport, and that resistant hosts limit the supply of copper to *S. sclerotiorum* by maintaining copper ion homeostasis. This research provides new insights into the interaction between *S. sclerotiorum* and the host, highlighting the importance of ROS and copper in these interactions.

## Results

### Copper is involved in the interaction between *Brassica oleracea* and *S. sclerotiorum*

*S. sclerotiorum* induces typical lesions, which are the main battlegrounds of gene interactions between *S. sclerotiorum* and the host. We previously detected differentially expressed genes (DEGs) by comparing gene expression in lesions of resistant and susceptible F_2_ plants of *B*. *oleracea* (Ding *et al*, 2019). Here, the set of transcriptome data was analyzed for dynamic changes of gene expression in sclerotinia and hosts during infection. A total of 738 and 228 *S. sclerotiorum* DEGs (24 hours post inoculation [hpi] vs 12 hpi) were detected in lesions of resistant and susceptible *B. oleracea*, respectively (Fig EV1A), which were significantly enriched for three overlapping Gene Ontology (GO) terms, ‘oxidation–reduction process’, ‘copper ion transport’ and ‘copper ion import’ (Fig 1A). Eight *S. sclerotiorum* DEGs involved in the ‘copper ion transport’ and ‘copper ion import’ processes were up-regulated during infection as revealed by both RNA-seq analysis and qRT-PCR analysis (Figs EV1B and Appendix Figure S1A).

**Figure 1.**
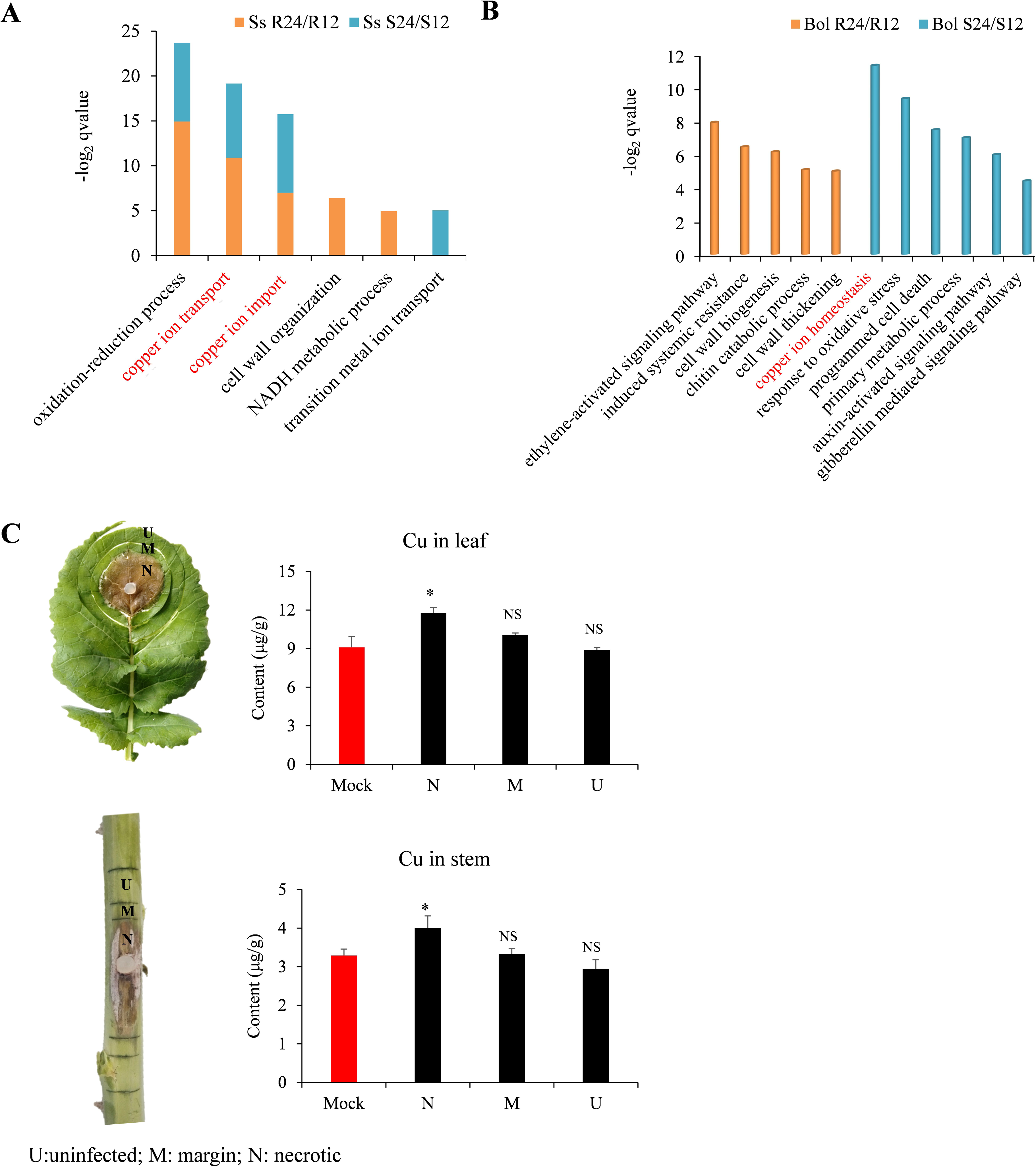
Copper is involved in the interaction between *Brassica* and *Sclerotinia sclerotiorum*. A Gene Ontology (GO) biological processes represented by *S. sclerotiorum* DEGs (differentially expressed genes; 24 hours post inoculation [hpi] vs 12 hpi) in infected *B. oleracea* stems. B GO biological processes represented by resistant and susceptible *B. oleracea* DEGs (24 hpi vs 12 hpi). C Copper concentration in uninfected (U), margin (M) and necrotic (N) tissues of *B. napus* leaves and stems at 48 hpi. Error bars indicate the standard deviation of three replicates. *: represents significant difference compared to the mock value at 0.05 level, NS indicates no significant difference at 0.05 level (Student’s *t*-test). Ss R24/R12: *S. sclerotiorum* DEGs in resistant *B. oleracea* identified by comparing 24 hpi to 12 hpi; Ss S24/S12: *S. sclerotiorum* DEGs in susceptible *B. oleracea* identified by comparing 24 hpi to 12 hpi; Bol R24/R12: *B. oleracea* DEGs in resistant *B. oleracea* identified by comparing 24 hpi to 12 hpi; Bol S24/S12: *B. oleracea* DEGs in susceptible *B. oleracea* identified by comparing 24 hpi to 12 hpi.

A total of 5988 and 5441 DEGs (24 hpi vs 12 hpi) were detected and subjected to GO analysis in resistant and susceptible *B. oleracea* stems, respectively (Fig EV1C). Interestingly, the biological process ‘copper ion homeostasis’ was significantly enriched in susceptible *B. oleracea* but not in resistant *B. oleracea* (Fig 1B). Among ten DEGs involved in ‘copper ion homeostasis’ (Fig EV1D), seven genes (*Bol023613*, *Bol026950*, *Bol044257*, *Bol002542*, *Bol011307*, *Bol000591* and *Bol029708*) with consistent expression patterns between RNA-seq analysis and qRT-PCR analysis were significantly down-regulated in susceptible *B. oleracea* plants, but only slightly down-regulated or stably expressed in resistant *B. oleracea* plants (Appendix Figure S1B). We further analyzed their expression in parental resistant (C01) and susceptible (C41) *B. oleracea* lines via qRT-PCR. All seven genes showed sharply down-regulated expression in susceptible parental line C41, while six of the seven genes (*Bol023613*, *Bol044257*, *Bol002542*, *Bol011307*, *Bol000591* and *Bol029708*) showed a stably or even slightly up-regulated expression in the resistant parental line C01 (24 hpi vs 12 hpi) (Appendix Figure S1C). This suggests that copper ion homeostasis may be disrupted in susceptible *B. oleracea* but not in resistant *B. oleracea* during early infection.

### Copper is transported into the necrotic area from healthy area during infection

We analyzed the copper distribution in and around lesions of infected leaves and stems in the moderately resistant rapeseed (*Brassica napus*) cultivar Zhongshuang 11 at 48 hpi. The content of copper in the necrotic areas (Cu_leaf-N_ = 11.76 μg/g, Cu_stem-N_ = 4.00 μg/g) was significantly higher than that in the uninfected (Cu_leaf-U_ = 8.90 μg/g, Cu_stem-U_ = 2.94 μg/g) and margin (Cu_leaf-Ma_ = 10.05 μg/g, Cu_stem-Ma_ = 3.33 μg/g) areas in both leaves and stems (*P* < 0.05), and significant differences in copper content were found in necrotic areas compared to mock-infected tissue (Cu_leaf-Mock_ = 9.09 μg/g, Cu_stem-Mock_ = 3.29 μg/g) (*P* < 0.05) (Fig 1C). These data indicate that copper is transported into the necrotic area from healthy area during early infection.

### Copper ion homeostasis genes promote host resistance

To test whether copper ion homeostasis is associated with host resistance, the six genes in this process that were stably expressed or slightly up-regulated in resistant *B. oleracea* but significantly down-regulated in susceptible *B. oleracea* were aligned with four *A. thaliana* orthologs (*AtCCS*, *AtMT2A*, *AtDRT112* and *AtCCH*) (Appendix Figure S2A). We tested the function of these *A. thaliana* homologs with respect to *S. sclerotiorum* resistance using T-DNA mutants (*atccs*, *atmt2a*, *atdrt112* and *atcch*) and overexpression lines (OX-AtCCS, OX-AtMT2A, OX-AtDRT112 and OX-AtCCH) (Appendix Figure S2B and C). Notably, all of the reduced-expression mutant lines were more susceptible to *S. sclerotiorum* compared to the wild type, while the overexpression lines displayed higher resistance (Fig 2A). At 24 hpi, the lesion size was 1.28–1.35 cm^2^ in the T-DNA mutant lines, 1.06–1.07 cm^2^ in the wild type and 0.59–0.62 cm^2^ in the overexpression lines (Fig 2B).

**Figure 2.**
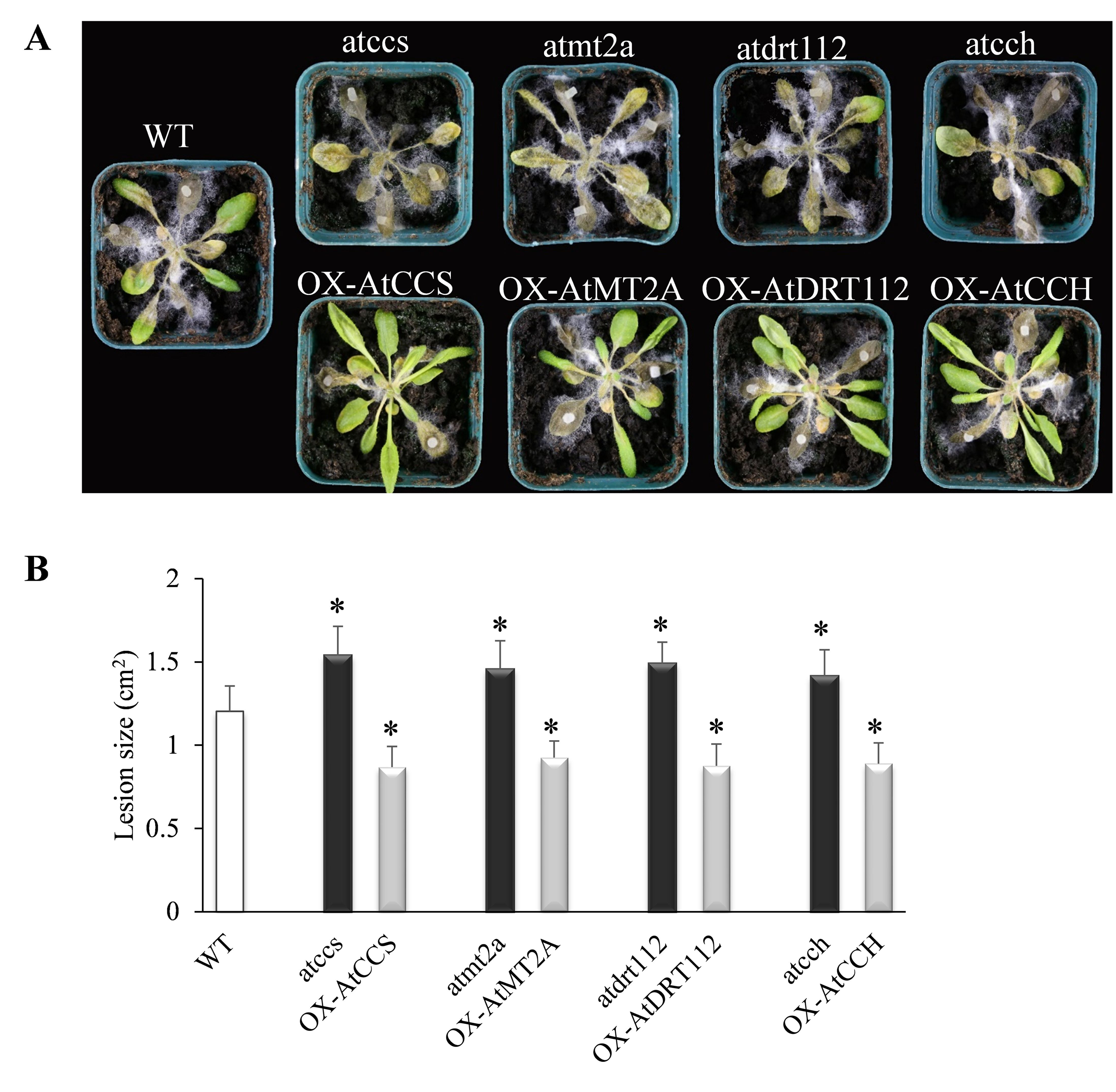
Copper ion homeostasis genes promote resistance to *S. sclerotiorum* in *Arabidopsis*. A Disease symptoms in *Arabidopsis* wild type (WT), T-DNA mutants (*atccs*, *atmt2a*, *atdrt112* and *atcch*) and overexpression lines (OX-AtCCS, OX-AtMT2A, OX-AtDRT112 and OX-AtCCH) corresponding to copper ion homeostasis genes at 4 days post inoculation (dpi) of *S. sclerotiorum* wild-type strain 1980. B Lesion size at 24 hpi in A. Error bars indicate the standard deviation of six replicates. *: represents significant difference from WT at 0.05 level (Student’s *t*-test).

### Copper ion homeostasis is associated with response to oxidative stress in the host

To further explore how copper ion homeostasis is associated with host resistance, the transcriptomes of leaves from *A. thaliana* wild type, T-DNA mutants and overexpression lines of *AtCCS*, *AtMT2A*, *AtDRT112* and *AtCCH* at 0, 6, and 12 hpi were sequenced, producing an average of 22.9 million clean reads for each sample. On average, 17.7 and 0.97 million clean reads were mapped to the reference genome of *A. thaliana* and *S. sclerotiorum* per sample, respectively. As expected, the overexpression lines exhibited higher expression of the corresponding target genes than the T-DNA mutants and the wild type line (Fig EV2A). Five common GO terms, including ‘response to oxidative stress’, were detected by comparing the up-regulated DEGs between overexpression lines and T-DNA mutants and between overexpression lines and wild type at 12 hpi (Fig 3A). Considering that more DEGs were found between the overexpression lines and T-DNA mutants than between the overexpression lines and wild type, we conducted a Weighted Gene Co-expression Network Analysis (WGCNA) using a total of 7321 *A. thaliana* DEGs between the overexpression lines and T-DNA mutants (Table EV1). This analysis produced 17 modules (groups of genes with similar expression pattern), of which one (shown in red) showed a negative correlation with lesion size (*r* = −0.45, *P* = 0.03) (Fig EV2B). It encompassed 394 genes, which were significantly enriched for the biological process of ‘response to oxidative stress’ (Fig 3B). In addition, total of 1273 *S. sclerotiorum* DEGs (overexpression lines vs. T-DNA mutants) were detected and used for WGCNA, which resulted in five significant modules (*P* < 0.05), two of which were highly correlated with peroxisome organization and oxidation–reduction (Table EV2; Fig EV2C).

**Figure 3.**
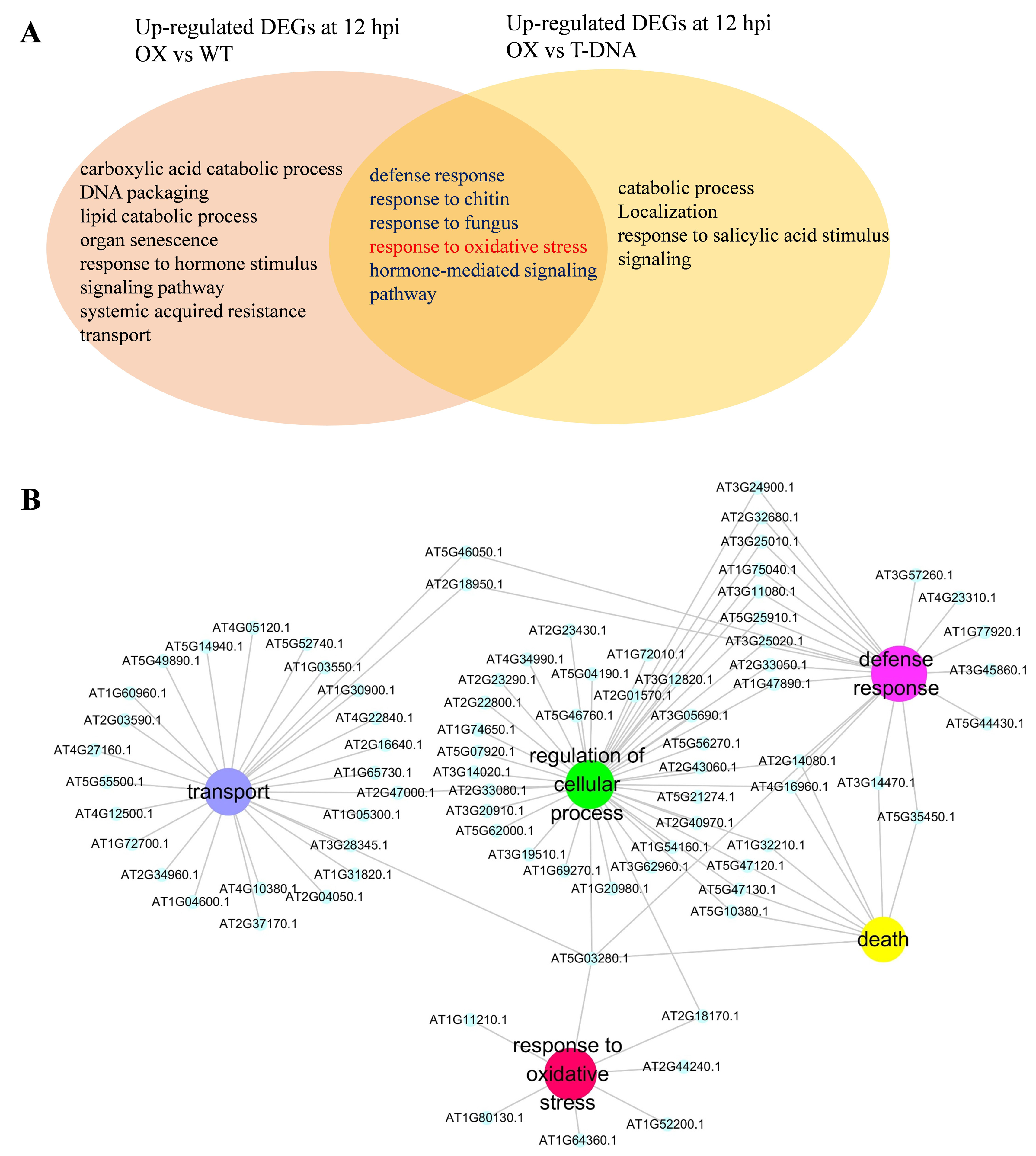
The biological process of ‘copper ion homeostasis’ is associated with ‘response to oxidative stress’ in the host. A GO terms (overlapped among the four genes) significantly enriched among the up-regulated DEGs at 12 hpi between *Arabidopsis* overexpression (OX) lines and wild type (WT) and between OX and T-DNA mutants. B GO terms significantly enriched among 394 DEGs in the module that significantly and negatively correlated with lesion size based on WGCNA (Weighted Gene Co-expression Network Analysis). The network was visualized using Cytoscape v3.4.

The oxidative burst is a typical response of a host against pathogen attack (Jones and Dangl, 2006). To functionally test whether copper ion homeostasis genes are associated with response to oxidative stress, we analyzed the antioxidant activity of these overexpression lines and T-DNA mutants by staining inoculated leaves at 0, 6 and 12 hpi with 3,0-diaminobenzidine (DAB) and nitrotetrazolium blue chloride (NBT) for the accumulation of H_2_O_2_ and ·O_2_^-^, respectively. We observed deeply stained areas around the inoculant column at both 6 and 12 hpi in the T-DNA mutant lines, but more lightly stained areas in the overexpression lines (Fig 4A), suggesting that antioxidant activity was enhanced in the overexpression lines but suppressed in the T-DNA mutants. This conclusion was further supported by a Cu/Zn SOD enzyme activity assay, in which the overexpression lines exhibited 41.79- to 45.93 and 49.62 to 66.97 U/mgprot, and T-DNA mutants exhibited 14.43 to 22.03 and 10.71 to 26.77 U/mgprot enzyme activity at 6 and 12 hpi, respectively (Fig 4B). Wild type line was in the middle with 32.14 and 42.82 U/mgprot at 6 and 12 hpi, respectively. These results indicate that copper ion homeostasis is associated with the detoxification of ROS in the host.

**Figure 4.**
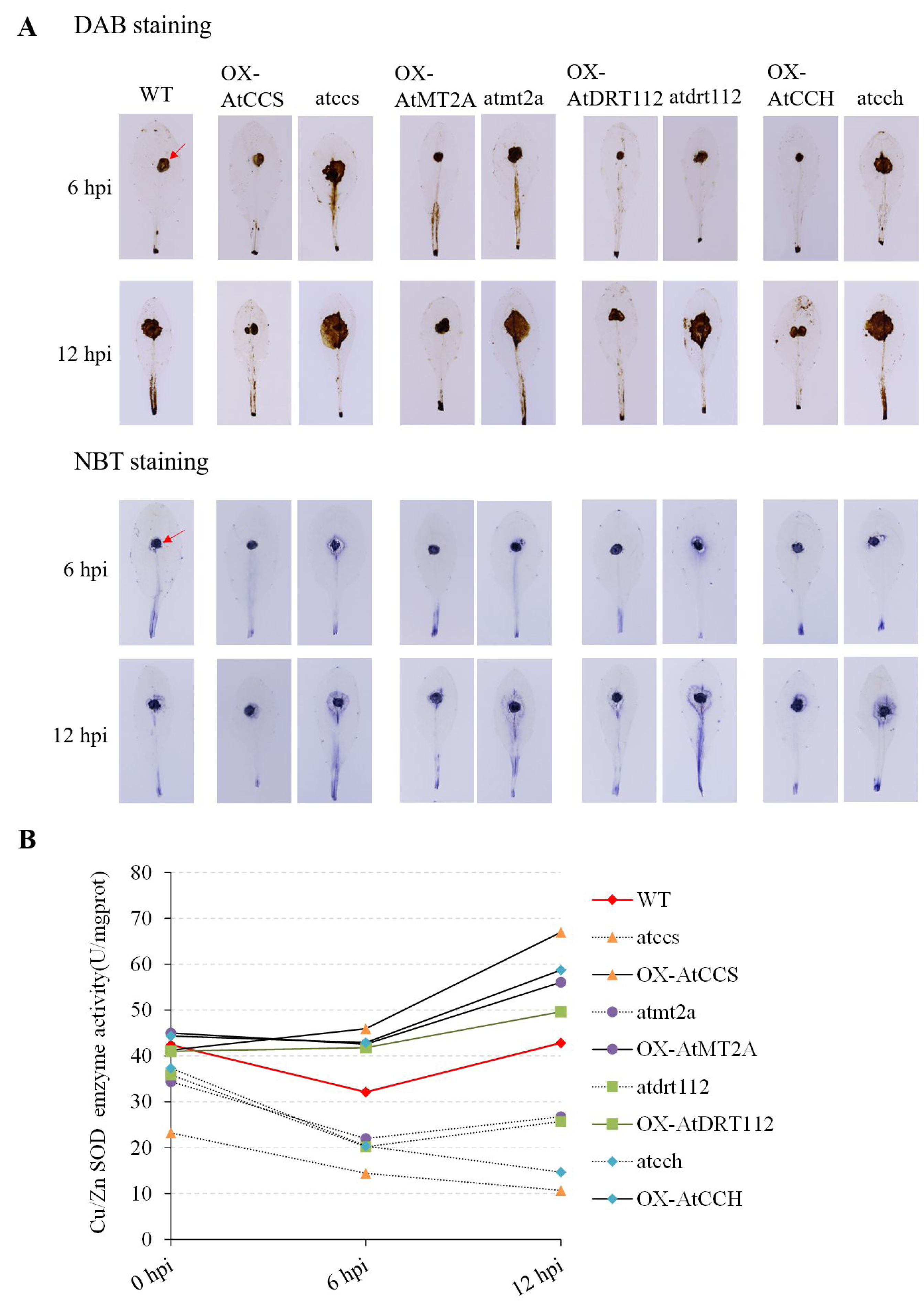
Copper ion homeostasis genes promote antioxidant activity in host. Antioxidant activity assays in wild type (WT), overexpression lines (OX) and T-DNA mutants of copper ion homeostasis genes in *Arabidopsis* at 0, 6 and 12 hpi. A DAB (H_2_O_2_ accumulation) and NBT (·O_2_^-^ accumulation) staining. One representative replicate from the five experiments is shown. The red arrows indicate the inoculant columns. B Enzyme activity of Cu/Zn SOD in leaves of *Arabidopsis* at 0, 6 and 12 hpi. Error bars indicate standard deviation for three replicates.

Considering that necrotrophic pathogens exploit the host oxidative burst, which contributes to pathogen attack (Heller and Tudzynski, 2011; Govrin and Levine, 2000), we hypothesized that copper ion homeostasis was associated with host resistance against diverse necrotrophic fungal pathogens. To test this hypothesis, we examined host resistance against another typical necrotrophic pathogen, *Botrytis cinerea* (Amselem *et al*, 2011). The *Arabidopsis AtCCS*, *AtMT2A*, *AtDRT112* and *AtCCH* overexpression lines increased resistance to *B. cinerea* by 30–60%, whereas the corresponding T-DNA mutants reduced resistance by 20–30% in comparison with wild type at 24 hpi (Fig EV3).

### *S. sclerotiorum* requires trace copper for pathogen infection

Copper is an essential nutrient for microbial pathogens and serves as an important cofactor of enzymes that scavenge ROS (Banci *et al*, 2012; Carr and Winge, 2003). To test the hypothesis that *S. sclerotiorum* takes up and utilizes host-derived copper ions for the synthesis or activation of enzymes involved in ROS scavenging during infection, we explored the roles of three *S. sclerotiorum* DEGs annotated as involved in ‘copper ion transport/import’: *SS1G_05578* (*SsCTR1*), *SS1G_00102* (*SsCCS*) and *SS1G_10888* (*SsATX1*) with their silenced and overexpressing strains (Fig EV4A and B). The products of these genes function to transport extracellular copper into the fungal cell (*SsCTR1*), deliver copper to Cu/Zn SOD (*SsCCS*) and detoxify the oxidative damage (*SsATX1*) (Cobine *et al*, 2006; Ding *et al*, 2011; Himelblau *et al*, 1998). These *S. sclerotiorum* genes exhibited down-regulated expression in all of the silenced strains (Sictr1, Siccs and Siatx1), and up-regulated expression in all of the overexpressing strains (OXctr1, OXccs and OXatx1) (Fig 5A), indicating they coordinately expressed in *S. sclerotiorum*. The lesion size after inoculation with the silenced strains was smaller than those after inoculation with the wild-type strain, while the largest lesion size was observed after inoculation with the overexpressing strains in the detached leaves of *B. napus* and *Arabidopsis* seedlings (Fig EV4C-E), indicating that these genes are involved in the virulence of *S. sclerotiorum*.

**Figure 5.**
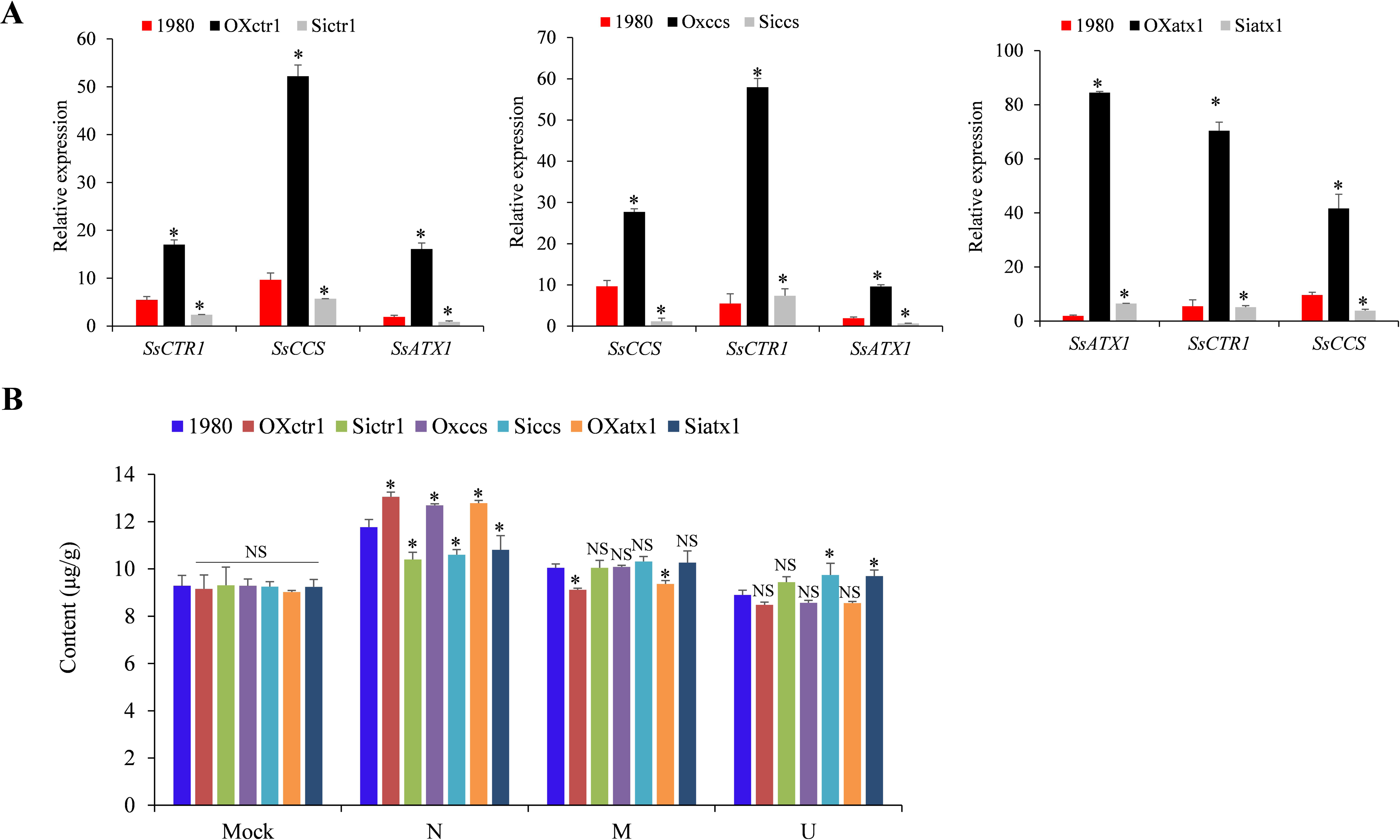
*SsCTR1*, *SsCCS* and *SsATX1* associated with copper absorption during infection. A Relative expression of *SsCTR1*, *SsCCS* and *SsATX1* in wild-type strain 1980, silenced and overexpressing strains as indicated by qRT-PCR analysis. The quantity of *S. sclerotiorum Tubulin* cDNA was used to normalize different samples. Error bars indicate the standard deviation of three independent samples. B Copper concentration in uninfected (U), margin (M) and necrotic (N) tissues of *B. napus* leaves at 48 hpi with wild-type, overexpressing and silenced strains. Error bars indicate the standard deviation of three replicates. *: represents significant difference between the transgenic strains and wild-type strain at the level of 0.05. NS indicates no significant difference at 0.05 level (Student’s *t*-test).

Copper content in the leaves infected by the silenced and overexpressing strains of these three genes was determined. All the strains showed higher copper content in the necrotic areas than the uninfected and margin tissues (Fig 5B). The content of copper in the necrotic areas of overexpressing strains contained 7.90%-10.93% more copper than that of wild-type, and the content of copper in the necrotic areas of silenced strains contained 11.62%-16.38% less copper than that of wild-type (Fig 5B). These results suggest that *SsCTR1*, *SsCCS* and *SsATX1* may associated with copper uptake in *S. sclerotiorum* during infection. To test the role of copper in the virulence of *S. sclerotiorum*, we sprayed low-concentration CuSO_4_ solutions (0.01, 0.025 and 0.05 mg/L) onto the leaf surface and incubated for 30 min prior to inoculation with each of the silenced strains. The lesion sizes after inoculation with the three silenced strains increased by 1.1- to 1.4-fold at 48 hpi in comparison with the non-CuSO_4_ treatment (sprayed with ddH_2_O), but exhibited no significant difference from the treatment that was sprayed with ddH_2_O and inoculated with the wild-type strain 1980 (Fig 6A and B). However, treatment with highly concentrated CuSO_4_ solutions (0.1 and 0.25 mg/L) significantly reduced disease symptoms of the silenced strains (*P* < 0.05) (Fig 6B). Similar observations were detected in the wild-type and overexpressing stains, which increased lesion size under low concentration of CuSO_4_, but decreased lesion size in high concentration of CuSO_4_ (Fig 6B). We further found that low-concentration CuSO_4_ promoted the growth of all the stains on potato dextrose agar (PDA) plates, and that high-concentration CuSO_4_ suppressed the growth of *S. sclerotiorum* (Fig 6C). Especially, the growth inhibition was higher in the silenced strains than the overexpressing strains in high-concentration CuSO_4_ (0.25 mg/L) (Fig 6C). These findings suggest that low-concentration copper could promote the growth of strains, and restores the virulence of the silenced strains. It indicates that trace copper is required for the growth and infection of *S. sclerotiotum*.

**Figure 6.**
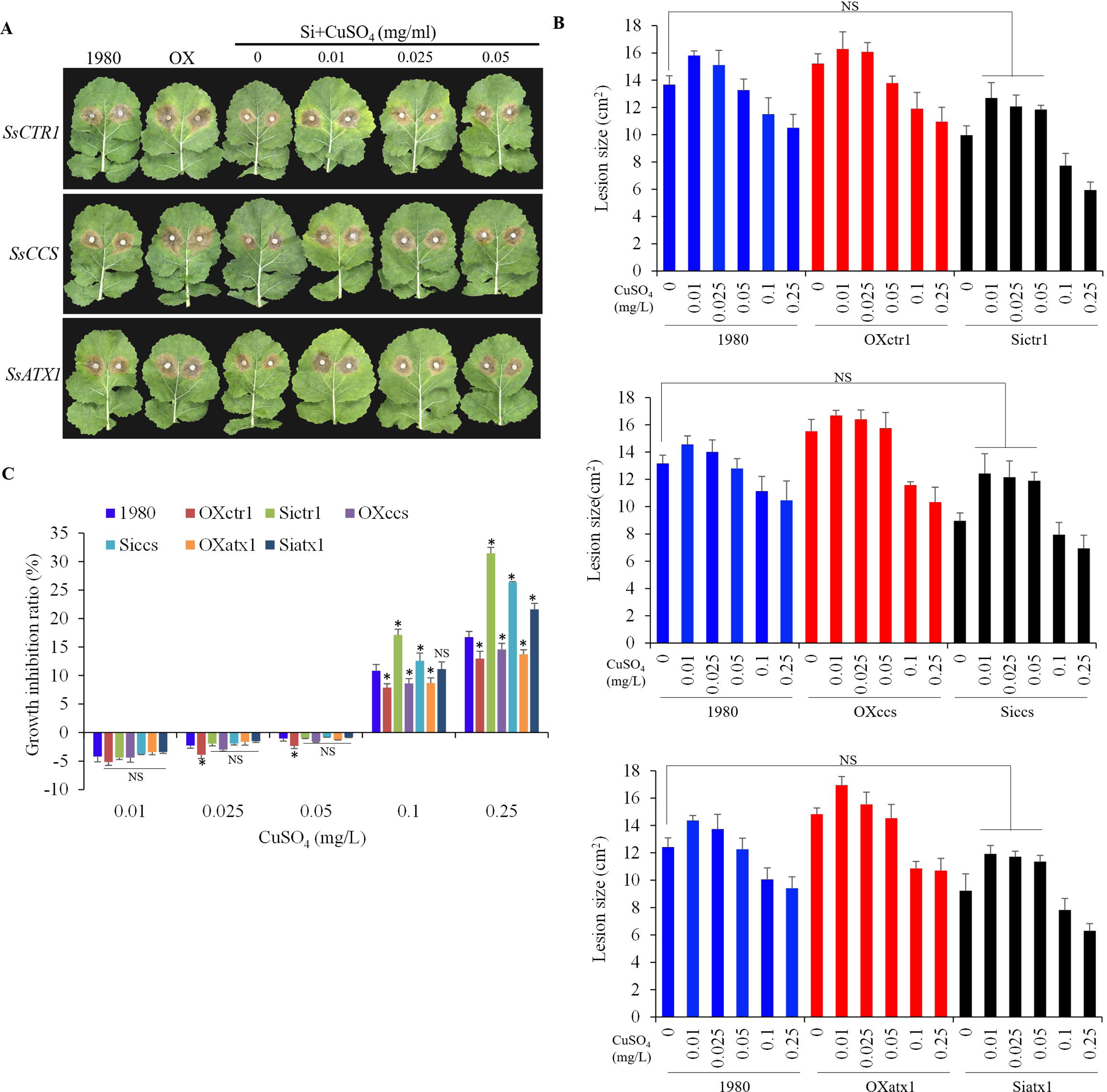
Trace copper restores the virulence of silenced strains of *S. sclerotiorum*. A Disease symptoms of leaves after inoculation with *S. sclerotiorum* wild-type strain 1980, overexpressing strains (OX, OXccs, OXatx1, OXcox17 and OXctr1) and silenced strains (Si, Siccs, Siatx1, Sicox17 and Sictr1) for genes involved in ‘copper ion transport/import’. The tests of the silenced strains were supplemented with low concentrations of CuSO_4_ solution, while the tests of 1980 and the overexpressing strains were sprayed with water. One representative replicate from the five experiments is shown. B Quantitation of lesion sizes at 48 hpi produced by wild-type, overexpressing and silenced strains when supplemented with low and high concentrations of CuSO_4_ solution. Error bars indicate standard deviation of five replicates. NS: represents no significant difference from wild-type strain at 0.05 level (Student’s *t*-test). C The growth on PDA for wild-type, overexpressing and silenced strains with supplementing low and high concentration of CuSO_4_. Error bars indicate standard deviation of five replicates. *: represents significant difference between the transgenic strains and wild-type strain at the level of 0.05. NS: represents no significant difference from wild-type strain at 0.05 level (Student’s *t*-test).

### Copper ion transport/import involves in ROS scavenging in *S. sclerotiorum*

To further test the hypothesis that *S. sclerotiorum* utilizes copper to promote ROS scavenging, we observed superoxide (·O_2_^-^) accumulation among wild-type strain 1980, the silenced strains and overexpressing strains of *SsCTR1*, *SsCCS* and *SsATX1* using NBT staining. The accumulation of ·O_2_ in the fungal hyphal tips was highest in the silenced strains, followed by the wild-type strain and the overexpressing strains (Fig 7A). We then cultured all strains on PDA supplemented with different concentrations of H_2_O_2_ (0, 2, 6, 10 and 15 mM) (Fig 7A). The growth of the silenced strains was most seriously inhibited (reduced by 26.2–100%), followed by the wild-type strain (reduced by 18.0–90.3%) and the overexpressing strains (reduced by 2.2–68.4%) (Fig 7B). These observations indicate that ROS can inhibit the growth of *S. sclerotiorum* and that these *S. sclerotiorum* genes are involved in the detoxification of ROS.

**Figure 7.**
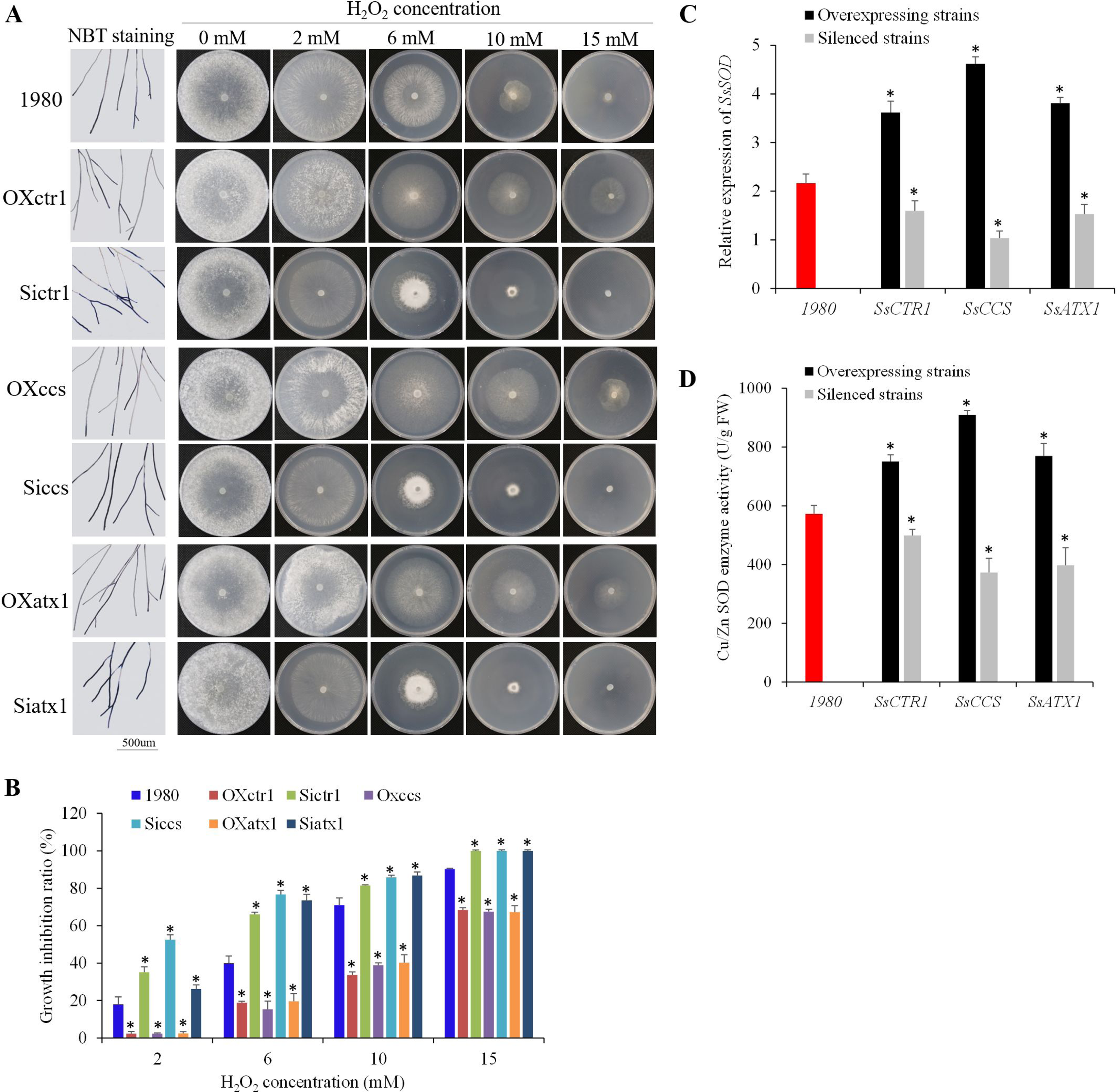
*SsCCS* promotes antioxidant activity in *S. sclerotiorum*. A Accumulation of ·O_2_^-^ (NBT staining at 24 hpi) in the hyphae tips and growth phenotypes on PDA supplemented with different concentrations of H_2_O_2_ at 4 dpi. One representative replicate from the five experiments is shown. B The inhibition rate of hyphal growth on PDA supplemented with different concentrations of H_2_O_2_. Error bars indicate standard deviation of five replicates. *: represents significant difference between the transgenic strains and wild-type strain at the level of 0.05. NS: represents no significant difference from wild-type strain at 0.05 level (Student’s *t*-test). C Relative expression of *S. sclerotiorum SsSOD1* in wild-type strain 1980, silenced and overexpressing strains as indicated by qRT-PCR analysis. The quantity of *S. sclerotiorum Tubulin* cDNA was used to normalize different samples. Error bars indicate the standard deviation of three independent samples. *: represents significant difference between the transgenic strains and wild-type strain at the level of 0.05 (Student’s *t*-test). D Enzyme activity of Cu/Zn SOD in hyphae of wild-type, silenced and overexpressing strains. Error bars indicate the standard deviation of three independent samples. *: represents significant difference between the transgenic strains and wild-type strain at the level of 0.05 (Student’s *t*-test).

Because Cu/Zn SOD is one of the primary superoxide scavengers (Mittler, 2017), we analyzed the *SsSOD1* expression level in the wild-type, silenced, and overexpressing strains. In the silenced strains, *SsSOD1* transcript accumulation was only 0.5–0.7 times the wild-type levels, while the overexpressing strains showed higher of *SsSOD1* expression, by 1.7- to 2.1-fold relative to wild-type strain (Fig 7C). In comparison with the wild-type (572.51 U/gFW), the Cu/Zn SOD enzyme activity was lower in silenced strains (372.88-499.39 U/gFW) and higher in overexpressing strains (750.44-909.86 U/gFW) (Fig 7D). These results indicate that these three *S. sclerotiorum* copper-related genes promote the expression of *SsSOD1* and increase enzyme activity of Cu/Zn SOD, resulting in the increase of ROS scavenging capacity in the fungal cells.

### Copper ion transport/import genes promote infection cushions formation and oxalic acid production in *S. sclerotiorum*

ROS is associated with the formation of infection cushions and production of oxalic acid in *S. sclerotiorum* (Liberti *et al*, 2013; Veluchamy *et al*, 2012; Kim *et al*, 2011). Infection cushions are essential for the *S. sclerotiorum* to penetrate the host cuticle and infect plants (Liang and Rollins, 2018), and oxalic acid (OA) is one of the virulence factors of *S. sclerotiorum* (Li *et al*, 2018; Liang and Rollins, 2018). Therefore, we monitored the infection cushions and OA production among the wild-type, silenced and overexpressing strains on *B. napus* leaves. Compared to the wild-type strain, the silenced strains exhibited fewer and looser infection cushions, while overexpressing strains exhibited more and compacter infection cushions (Fig EV5A), indicating that these *S. sclerotiorum* genes affect the establishment of infection.

Meanwhile, we detected significant differences in OA concentration among the wild-type, silenced and overexpressing strains after culturing for 2 days on PDA medium. The average OA concentration in the silenced strains was 6.30 mg/ml, followed by the wild-type (7.81 mg/ml) and overexpressing strains (9.39 mg/ml) (Fig EV5B). At the same time, the average pH value of the culture medium from the overexpressing strains (pH = 2.79) was lowest, followed by the wild-type (pH = 2.92) and silenced strains (pH = 3.13) (Fig EV7C). The OA synthesis gene *SsOAH* exhibited lower expression in the silenced strains than in the wild-type strain, but higher expression in the overexpressing strains (Fig EV5D). It suggests that expression disruption of the copper ion transport/import genes suppresses the OA production in *S. sclerotiorum*.

## Discussion

Copper is a component of numerous enzymes and plays a key role in the responses to oxidative stress (Culotta *et al*, 2006; Berterame *et al*, 2018). By analyzing differentially expressed genes (DEGs) at lesion for dynamic changes in host and *S. sclerotiorum* during infection, we found that the genes in the ‘copper ion import’ and ‘copper ion transport’ with up-regulated expression involved in *S. sclerotiorum* copper uptake, virulence, ROS detoxification, fungal establishment and OA production, and that the host genes in the ‘copper ion homeostasis’ with stable expression in the resistant line, but down-regulated expression in the susceptible line were associated with response to oxidative stress and resistance to *S. sclerotiorum*. Our data indicate a battlefield at the host-*S. sclerotiorum* interface where *S. sclerotiorum* hijacks host-derived copper to detoxify ROS in its cell, while the resistant host maintains ‘copper ion homeostasis’ to limit the supply of copper to the pathogen.

The generation of ROS at the infection site is one of the earliest responses of pathogen-associated molecular-pattern-triggered immunity (PTI) (Jones and Dangl, 2006). As secondary messengers, ROS are indispensable for signaling, stress responses and developmental processes (Marschall and Tudzynsk, 2014). However, excess ROS trigger programmed cell death (PCD) and cause host necrosis, which facilitates the growth of necrotrophic pathogens (Heller and Tudzynski, 2011; Kim *et al*, 2008; Lu and Higgins, 1999). Antioxidant components in the host, such as peroxidase, SODs, glutathione sulfhydryl transferase (GST) and glutathione (GSH) are associated with resistance against *S. sclerotiorum* (Liang *et al*, 2008; Yang *et al*, 2007; Wen *et al*, 2013; Wei *et al*, 2016; Mei *et al*, 2016; Garg *et al*, 2013), and resistance against *S. sclerotiorum* can be improved by decreasing the accumulation and/or production of ROS in the host (Ranjan *et al*, 2018; Wang *et al*, 2014). The four DEGs in the biological process of ‘copper ion homeostasis’ we detected during infection are involved in various antioxidant activities. *AtCCS* is responsible for the activation of Cu/Zn SOD (Chu *et al*, 2005). Metallothioneins (MTs), which act as heavy metal chelators and ROS scavengers, contribute to plant adaptation to abiotic stresses (Kim and Kang, 2018). *DRT112* is one of two *Arabidopsis* plastocyanin genes (*PETE2*), which function to buffer excess copper (Abdel-Ghany, 2009). *AtCCH* is a homolog of yeast *ATX1*, which functions to deliver copper into laccase or Fet3 (Himelblau *et al*, 1998). In this study, we found that the overexpression of these genes in *Arabidopsis* enhanced host ROS detoxification and resistance against *S. sclerotiorum*, and the genes in the biological process of ‘copper ion homeostasis’ were coordinately expressed with those for ‘response to oxidative stress’. Therefore, our research provides evidence that copper ion homeostasis is associated with ROS detoxification in host.

Free copper ions induce ROS production (Rae *et al*, 1999), which may be toxic to *S. sclerotiorum*. We found that highly concentrated CuSO_4_ solutions suppressed the growth of *S. sclerotiorum*. In fact, copper is one of important active ingredients in many bactericidal and fungicidal agents, such as Bordeaux mixture (Martins *et al*, 2014). However, *S. sclerotiorum* can grow in necrotic areas with a relatively high concentration of copper. Three aspects of our findings might help explain this phenomenon. (1) The copper uptake system of *S. sclerotiorum* was elevated during infection. *SsCTR1*, which functions to import extracellular copper into the fungal cells (Samanovic *et al*, 2012), exhibited up-regulated expression, indicating that host-derived copper may be imported into the *S. sclerotiorum* cells. (2) A few genes associated with copper detoxification, such as MTs (*SsMT*), Sur7 (*SsSUR7*) and P-type ATPase (*SsATP7A*) (Weissman *et al*, 2000; Douglas *et al*, 2012; Ladomersky and Petris, 2015), were not significantly induced in *S. sclerotiorum* during infection (Appendix Figure S3), indicating that the amount of host-derived copper was not high enough to produce toxic effects in *S. sclerotiorum*. (3) *SsCCS*, *SsCOX17* and *SsATX1* expression was up-regulated during infection. The homologs of *SsCCS* function to deliver copper to oxidative scavengers Cu/Zn SOD (Gleason *et al*, 2014; Cobine *et al*, 2006). The ATX1 gene could act as a multi-copy suppressor of oxidative damage in yeast (Himelblau *et al*, 1998). It indicated that the copper might be utilized for the synthesis or activation of these ROS detoxification enzymes of *S. sclerotiorum* during infection. Thus, this study provides important insights into the question about survival of *S. sclerotiorum* at relatively high levels of ROS.

The idea of ‘nutritional immunity’ was first proposed to describe the resistance mechanism in human cells wherein they withhold transition metals, such as iron, zinc, manganese and copper to defend against microbial pathogen invaders (Besold *et al*, 2016; Malavia *et al*, 2017; Hood and Skaar, 2012; Crawford and Wilson, 2015). To overcome this strategy, successful pathogenic species must evolve specialized mechanisms to adapt to the nutritionally restrictive environment of the host and cause disease. For example, the human kidney and brain can thwart *Candida albicans* growth by limiting the supply of copper nutrients, producing ‘copper starvation’ to *C. albicans* (Besold *et al*, 2016). In response, *C. albicans* induces copper uptake machineries that enable it to survive in a copper-starved environment (Li *et al*, 2015). Here, the defense response in resistant lines that stabilized the expression of copper ion homeostasis genes and limited the availability of copper to *S. sclerotiorum* may be considered a form of nutritional immunity. To the best of our knowledge, this is the first description of nutritional immunity in plants. We therefore propose a possible host–*S. sclerotiorum* interaction model in which resistant plants induce nutritional immunity and restrict the supply of essential copper nutrients to *S. sclerotiorum* by maintaining copper ion homeostasis, while *S. sclerotiorum* enhances its copper uptake system and hijacks host-derived copper, which activates its ROS scavenging system during infection and promotes its survival and virulence (Fig. 8).

**Figure 8.**
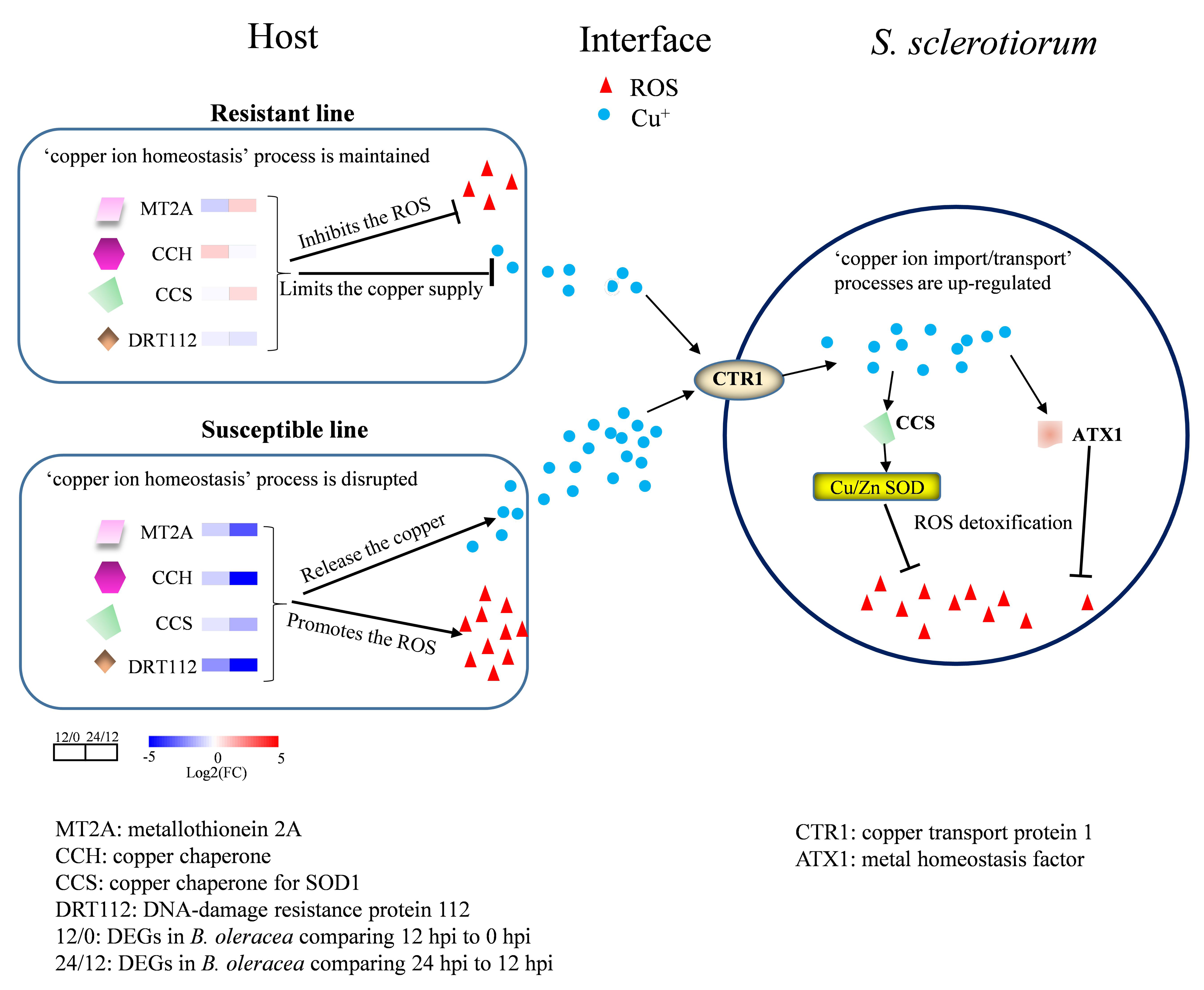
Model depicting the battle for copper acquisition between host and *S. sclerotiorum* during infection. *S. sclerotiorum* promotes its copper uptake systems which hijack host-derived copper, and activates its ROS scavenging system for survival and virulence, while a resistant host induces ‘nutritional immunity’ that restricts the supply of essential copper nutrients to *S. sclerotiorum* by maintaining ‘copper ion homeostasis’. 12/0: gene expression comparison of 12 hpi to 0 hpi in host, 24/12: gene expression comparison of 24 hpi to 12 hpi in host.

## Methods and materials

### Experimental strains and plants

The wild-type strain of *S. sclerotiorum* 1980 and *B. cinerea* wild-type strain B05.10 (Amselem *et al*, 2011) were used in this study. Fungal strains were grown on potato dextrose agar (PDA, 20% potato, 2% dextrose and 1.5% agar) at 22℃. *S. sclerotiorum* transformants were cultured on PDA amended with 80 μg/ml hygromycin B (Calbiochem, San Diego, CA) to stabilize the transformants. Seedlings from *A. thaliana* Col-0 (ecotype Columbia-0), T-DNA mutants and overexpression transgenic lines were grown in the autoclaved soil (Pindstrup) at 20 ± 2°C under a 12 h light/dark cycle with 70% relative humidity. The rapeseed cultivar Zhongshuang 11 was used for virulence assay.

### Vector construction and transformation of *S. sclerotiorum* and *A. thaliana*

The fragments of four *S. sclerotiorum* genes (*SsATX1*: 221 bp, *SsCOX17*: 232 bp, *SsCCS*: 221 bp and *SsCTR1*: 461 bp) were PCR-amplified from the cDNA library of *S. sclerotiorum* wild-type strain 1980 by using the specific primers of RNAi vector construction in Table EV3. The sense and antisense fragments were ligated into the plasmid vector pCIT (Yu *et al*, 2017) at the corresponding sites, and a hygromycin resistance gene cassette from pSKH (Hamid *et al*, 2013) was isolated and ligated into them, resulting the RNAi vectors, pSiatx, pSicox17, pSiccs and pSictr1. To generate *S. sclerotiorum* overexpression strains of these four genes, the full-length coding sequences were amplified by using the specific primers of overexpression vector construction in Table S1 and cloned into a hygromycin resistance containing vector, driven by the *S. sclerotiorum* constitutive expression promoter EF1-A (*SS1G_06124*, translation elongation factor 1 alpha), resulting the overexpression constructs, pOEATX1, pOECOX17, pOECCS and pOECTR1. *S. sclerotiorum* transformations were conducted with a standard polyethylene glycol (PEG)-mediated transformation protocol (Rollins, 2003).

The T-DNA mutant lines of *A. thaliana* (*AtCCS*: SALK_025986C*, AtCCH*: SALK_118605C*, AtMT2A*: SALK_021037C, and *AtDRT112*: SALK_135199C) were obtained from the Arabidopsis Biological Resource Center, Ohio State University, USA. The homozygous T-DNA insertion lines were confirmed with primers flanking the T-DNA insertions (Table EV3) and the left border primer LB1.3 (ATTTTGCCGATTTCGGAAC). To generate the overexpression lines, the coding sequences of these four genes were amplified from a cDNA library of *A. thaliana* leaves (Col [Columbia]-0). The amplicons were digested with *Xba*I and *Xho*I and ligated into the plant expression vector pBinGlyRed3, which contains a gene encoded red fluorescent protein (DsRed). The resulting vectors (pBinGlyRed-*AtCCS*, pBinGlyRed-*AtCCH*, pBinGlyRed-*AtMT2A* and pBinGlyRed-*AtDRT112*) were introduced into the *Agrobacterium tumefaciens* strain GV3101 by electroporation (Wise *et al*, 2006) and transformed into *A. thaliana* Col-0 by the floral dip method (Clough and Bent, 1998).

### RNA-seq and network analysis

In our previous study (Ding *et al*, 2019), the stems of resistant and susceptible *B. oleracea* plants in a F_2_ population which derived from the cross between a resistant *B. oleracea* genotype ‘C01’ (*B. incana*) and a susceptible *B. oleracea* genotype ‘C41’ (*B. oleracea* var. *alboglabra*) were inoculated with *S. sclerotiorum*, and lesions at 0, 12 and 24 hpi were collected for transcriptome sequencing. To analyze the dynamic changes during infection, here we analyzed the DEGs of *S. sclerotiorum* and *B. oleracea* by using the DESeq package (Anders and Huber, 2010).

To reveal the pathways associated with ‘copper ion homeostasis’, the RNA of infected leaves was sequenced from *A. thaliana* wild type (Col-0), T-DNA mutants and overexpression lines during infection. Briefly, the detached leaves were inoculated with *S. sclerotiorum* wild-type strain 1980 (Mei *et al*, 2016), and RNA from the lesions was extracted with the RNAprep pure Plant Kit (DP 432, Tiangen Biotech (BEIJING) CO., LTD). The sequencing library was generated using the Illumina RNA Library Prep Kit (NEB, USA) following the manufacturer’s recommendation, and sequenced on an Illumina HiSeq 4000 platform with three biological replicates. After removing low-quality reads and those with adapter sequences, poly-N sequence from the raw data, the clean reads were screened and aligned to the reference genomes of *A. thaliana* (https://www.arabidopsis.org/download/index.jsp) and *S. sclerotiorum* (http://fungidb.org/common/downloads/Current_Release/Ssclerotiorum1980UF-70/) by using the TopHat program (http://ccb.jhu.edu/software/tophat/index.shtml) (Trapnell *et al*, 2009) with default parameters except that the Q value was set to 100. Gene expression was quantified using htseq-count 0.6.1p2 (https://htseq.readthedocs.io/). The raw counts were normalized by TMM normalization using the edgeR package (Robinson *et al*, 2010) and the differential expression analysis was carried out using the DESeq package (Anders and Huber, 2010).

The threshold determining the significance of DEGs among multiple tests was set at a false discovery rate (*FDR*) ≤ 0.001 and |log_2_ ratio|≥ 1. GO (Gene Ontology) and KEGG (Kyoto Encyclopedia of Genes and Genomes) enrichment analyses were performed with an FDR ≤ 0.05 as the threshold using AgriGO (Tian *et al*, 2017) and KOBAS 3.0 (http://kobas.cbi.pku.edu.cn/), respectively.

Weighted correlation networks were produced among the DEGs with R package WGCNA (Weighted Gene Co-expression Network Analysis) (Langfelder and Horvath, 2008). Networks were visualized by Cytoscape v3.4 (Shannon *et al*, 2003).

### Measurement of copper ion concentration

Detached leaves and stems of *B. napus* cv. Zhongshuang 11, which was recognized as a moderately resistant rapeseed cultivar (Sun *et al*, 2017), were inoculated with *S. sclerotiorum* strains, and the uninfected (U), margin (M) and necrotic (N) tissues were collected at 48 hpi. The tissues were washed with distilled water, dried for 1 week at 80°C, and then washed with 11 N HNO_3_. Copper concentration of tissues was measured using atomic absorption spectroscopy (SPECTR AA220) at a wavelength of 324.8 nm.

### Quantitative RT-PCR

Gene expression was analyzed by qRT-PCR using a Bio-Rad CFX96 Real Time System (Bio-Rad, USA) and QuantiTect SYBR Green PCR master mix (Bio-Rad, USA), according to the manufacturer’s instructions. The *SsTubulin* and *BoActin3* genes were used as the internal control for *S. sclerotiorum* and *B. oleracea,* respectively. All the qRT-PCR primers were listed in Table EV3. The PCR cycling conditions comprised 1 cycle of 95°C for 30 s, then 39 cycles of 95°C for 5 s and 55-70°C for 1 min, followed by a melting curve ramping from 65°C to 95°C with temperature increasing by 0.5°C every 5 s (1 cycle). Transcript levels were calculated from the threshold cycle using the 2^-ΔΔCT^ method (Livak and Schmittgen, 2001). Three replicates were performed for each gene and data were analyzed using CFX Manager™ v3.0.

### Pathogenicity assays

Pathogenicity of *S. sclerotiorum* was evaluated by infecting *B. napus* and *A. thaliana* according to the procedure described previously (Ding *et al*, 2019; Mei *et al*, 2016). The detached leaves at seedling stage and stems at flowering stage of *B. napus*, and in vivo leaves of 3-week-old *A. thaliana*, were inoculated with mycelium-colonized agar plugs (0.6 cm for *B. napus*, 0.2 cm for *A. thaliana*) obtained from expanding margins of PDA-cultured colonies, with five replicates. The inoculation chamber was maintained at 85% relative humidity at 22°C. The lesion size (*S*, cm^2^) for the leaves was calculated with the formula *S* = π_*_*a*_*_*b*/4, where *a* and *b* represent the long and short diameter of an approximately elliptical lesion. The infection cushions on *B. napus* leaves at 6, 9 and 12 hpi were observed with a scanning electron microscope (JEOL JEM-6390LV).

### Antioxidant activity assays

Superoxide (·O_2_^-^) accumulation was assayed by staining wild-type and transformant hyphae of *S. sclerotiorum* with NBT (Kumar *et al*, 2014). Hyphae on the PDA plate at 24 hpi were infiltrated under gentle vacuum with NBT staining solution for 5 hours and then washed 3 times with distilled water, prior to observation with a microscope. Meanwhile, expression of the *S. sclerotiorum* Cu/Zn SOD gene (*SsSOD1*) was assayed by qRT-PCR, and the growth inhibition ratio was calculated by measuring the diameter every 12 hours when cultured on PDA in the presence of 2, 6, 10 and 15 mM H_2_O_2_. The enzyme activity of Cu/Zn SOD of hyphae was tested using an enzyme activity kit (A001-1; Nanjing Jiancheng Bioengineering Institute) according to the manufacturer’s protocol.

The antioxidant activity of *Arabidopsis* during infection was determined by DAB and NBT staining (Kumar *et al*, 2014). Leaves were infiltrated under gentle vacuum with DAB or NBT staining solution for 5 hours. After staining, the staining solution was replaced with bleaching solution (ethanol: acetic acid: glycerol = 3:1:1). After 15 ± 5 min in a boiling water bath (∼90-95 °C), the bleaching solution was replaced with fresh bleaching solution and stained in 60% glycerin. Meanwhile, the enzyme activity of Cu/Zn SOD was tested at 0, 6 and 12 hpi using an enzyme activity kit (A001-1; Nanjing Jiancheng Bioengineering Institute) according to the manufacturer’s protocol.

### Oxalic acid assays

Five agar plugs (6 mm) from the advancing edge of each *S. sclerotiorum* strain were transferred into 5 ml PDA medium with 50 mg/L bromophenol blue and incubated at 20°C with shaking at 150 rpm for 2 days. Prior to assaying the concentration of OA in the solution, a standard curve was generated using OA standard samples with a spectrometer (Evolution™ 201/220, ThermoFisher, USA). Meanwhile, the pH value of the solution was measured with a pH meter (INESA, China), and the expression of *SsOAH* was assayed in *S. sclerotiorum* strains by qRT-PCR.

## Acknowledgements

This study was financially supported by the National Natural Science Foundation of China (31671726, 31801395 and 31971978), the Natural Science Foundation of Chongqing (cstc2017shms-xdny80050, cstc2019jcyj-zdxmX0012 and cstc2019jcyj-msxm2511) and Fundamental Research Funds for the Central Universities (XDJK2018AA004 and XDJK2018B022). We sincerely acknowledge Dr. Pradeep Kachroo and Dr. Zhonglin Mou for critical comments.

## Author contributions

W.Q. designed the experiments, Y.D., J.M. performed experiments, analyzed the data, and wrote the manuscript. Y.C. performed and analyzed the RNA sequencing experiments. W.Y. performed cloning and transformation experiments. Y.M. performed copper ion concentration measurement and oxalic acid assays. B.Y., Y.Y. and K.R. performed the pathogenicity and antioxidant activity analysis. J.O.D and J. L. analyzed the data and helped write the manuscript. All authors reviewed the manuscript before publication.

## Expanded View Figure legends

**Figure EV1. DEG (differentially expressed gene) analysis of *Sclerotinia sclerotiorum* and *Brassica oleracea* (24 hours post inoculation [hpi] vs 12 hpi).**

A DEGs of *S. sclerotiorum* during infection in the resistant (R-Ss) and susceptible (S-Ss) *B. oleracea*. B DEGs of resistant (R-Bol) and susceptible (S-Bol) *B. oleracea*.

C Heat map of *S. sclerotiorum* DEGs involved in the process ‘copper ion import’ and ‘copper ion transport’.

D Heat map of *B. oleracea* DEGs involved in the process ‘copper ion homeostasis’. Ss R24/R12: the *S. sclerotiorum* DEGs in resistant *B. oleracea* by comparing 24 hpi to 12 hpi; Ss S24/S12: the *S. sclerotiorum* DEGs in susceptible *B. oleracea* by comparing 24 hpi to 12 hpi; Bol R24/R12: the *B. oleracea* DEGs in resistant *B. oleracea* by comparing 24 hpi to 12 hpi; Bol S24/S12: the *B. oleracea* DEGs in susceptible *B. oleracea* by comparing 24 hpi to 12 hpi.

**Figure EV2. DEGs and DEG network analysis of *Arabidopsis* and *S. sclerotiorum* during infection.**

A Relative expression level of target genes in the *Arabidopsis* T-DNA mutants and overexpression lines (OX) in comparison with the wild type Col-0 as revealed by the RNA-seq.

B Weighted Gene Co-expression Network Analysis (WGCNA) of the DEGs between OX lines and T-DNA mutants in *Arabidopsis*.

C WGCNA of *S. sclerotiorum* DEGs during infection of *Arabidopsis* overexpression lines and T-DNA mutants.

**Figure EV3. Copper ion homeostasis-related genes in *Arabidopsis* are associated with resistance to *Botrytis cinerea*.**

A Disease symptoms at 24 hpi in *Arabidopsis* wild type (WT), T-DNA mutants (atccs, atmt2a, atdrt112 and atcch) and overexpression lines (OX-AtCCS, OX-AtMT2A, OX-AtDRT112 and OX-AtCCH) after inoculation of *B. cinerea* strain B05.10.

B Lesion size in (A). Error bars indicate the standard deviation of six replicates. *: represents significant difference of WT with T-DNA mutants and overexpression lines of *Arabidopsis* at the level of 0.05 (Student’s *t*-test).

**Figure EV4. Virulence assays of the wild-type, silenced and overexpressing strains of *S. sclerotiorum*.**

A The silenced (RNAi: RNA interference) and overexpressed (OX) vectors.

B Relative expression level of the target genes in silenced, overexpressed and wild type strain 1980 on PDA medium as determined by qRT-PCR. The quantity of *S. sclerotiorum Tubulin* cDNA normalized different samples. Error bars indicate the standard deviation of three independent samples. *: represents significant difference between the transgenic strains and wild-type strain at the level of 0.05 (Student’s *t*-test).

C Virulence on *Brassica napus* Zhongshuang 11 in detached leaves at 48 hpi. Error bars indicate the standard deviation of six replicates. *: represents significant difference between the transgenic strains and wild-type strain at the level of 0.05 (Student’s *t*-test).

D Virulence in *Arabidopsis thaliana* plants at 24 hpi. Error bars indicate the standard deviation of six replicates. *: represents significant difference between the transgenic strains and wild-type strain at the level of 0.05 (Student’s *t*-test).

E Disease symptoms at 4 dpi (days-post inoculation) in *A. thaliana* plants.

**Figure EV5. The expression disruption of the copper ion transport/import genes suppresses the infection cushion formation and OA production in *S. sclerotiorum*.**

A Infection cushion observation on *B. napus* leaves inoculated with the wild-type, silenced and overexpressing strains at 6, 9 and 12 hpi. Bar = 100 μm. Experiments were repeated with three times with similar results.

B OA production of *S. sclerotiorum* strains. Error bars indicate the standard deviation of three replicates.

*: represents significant difference between the transgenic strains and wild-type strain at the level of 0.05 (Student’s *t*-test).

C Ambient pH of strains in PDA.

D The relative expression of *S. sclerotiorum SsOAH* among *S. sclerotiorum* strains. The quantity of *S. sclerotiorum Tubulin* cDNA normalized different samples. Error bars indicate the standard deviation of three independent samples. *: represents significant difference between the transgenic strains and wild-type strain at the level of 0.05 (Student’s *t*-test).

## Expanded View Table legends

**Table EV1.** 7321 *Arabidopsis* DEGs between the overexpression lines and T-DNA mutants.

**Table EV2.** 1273 *S. sclerotiorum* DEGs during infecting *Arabidopsis* overexpression lines and T-DNA mutants.

**Table EV3.** Primers used in this study.

